# Flow constraints at infection site shape multiplication-dissemination trade-offs and opposite regulatory programs of *Xanthomonas* and *Ralstonia* xylem pathogens

**DOI:** 10.64898/2026.02.17.706415

**Authors:** Amélie Caddeo, Matthieu Barret, Rémi Peyraud

## Abstract

Pathogens rely on multiple pathogenicity traits, such as proliferation, adhesion, motility and the production of virulence factors, to successfully colonize their host. The expression of virulence functions is often finely regulated to mitigate resource allocation trade-offs due to their cost for the cell. The ways in which constraints encountered inside the host shape strategies for mitigating the trade-off between pathogenicity traits remain poorly understood. *Xanthomonas campestris pv. campestris* (*Xcc*) and *Ralstonia solanacearum* (*Rs*) are two bacterial phytopathogens that can colonize the same plant habitat, the xylem vessel, from two different infection sites, leaf and root, respectively. Analyses of the virulent regulatory networks (VRN) of *Xcc* and *Rs* revealed differences in the expression of their virulence programs in a cell density-dependent manner. Notably, swimming motility and exopolysaccharide production were regulated using opposite strategies. Simulation of bacterial dispersion in the vascular system through spatial model showed that these VRNs were adapted to the constraint of xylem sap flow. This study reports how strong environmental constraints, such as the direction of xylem sap flow, can shape opposite regulatory programs and strategies for mitigating trade-offs for pathogens colonizing the same ecological niche.

**Importance:** The regulation of pathogen virulence programs and how differences in their execution confer invasion advantages remain poorly understood. Because the expression of pathogenicity traits is costly in terms of energy, pathogens must carefully control their deployment to proliferate efficiently within the host. This work uses a systems-level approach to provide a comprehensive understanding of the sequential expression of pathogenicity traits in two xylem pathogens *Ralstonia solanacearum* (*Rs*) and *Xanthomonas campestris pv. campestris* (*Xcc*) during xylem colonization. Moreover, computational analysis of their dispersal within xylem vessels highlights the significance of strategies that mitigate trade-offs to ensure effective colonization, despite the strong physical constraints imposed by xylem sap flow. Together, these results provide new insight into how distinct regulatory strategies influence the success of pathogen invasion and enhance our understanding of bacterial adaptation to host vascular environments.

## Introduction

Many bacterial pathogens, whether plant- or animal-related, use similar traits to infect their host and cause disease. These traits may include motility, adhesion, biofilm formation, exopolysaccharides (EPS) secretion, evasion from host immune system, and secretion of degrading enzymes (Reverchon et al. 2016; Piewngam et Otto 2024; Lu et al. 2025; Clements et al. 2012). The simultaneous expression of all these traits would be too costly in terms of energy for the cell to support division and multiplication (Ruparell et al. 2016; Diard et Hardt 2017; Sturm et al. 2011; Davis 2023). Hence, these traits can have a negative impact on each other due to a trade-off in resource allocation (Peyraud et al. 2018) or due to design constraints that are inherent to enzyme efficiency (Ferenci 2016). Moreover, activation should occur at appropriate timing to confer advantages during infection (Ramamurthy et al. 2020). Therefore, precise regulation and execution of the activation of pathogenicity traits, *i.e*. the regulatory program, is crucial for pathogens to successfully infect their host by mitigating the trade-off between all pathogenicity traits. However, it remains a limited understanding of how the regulatory program of a pathogen adapts to the complex spatial and temporal constraints encountered inside the host niches and how it mitigates the trade-off between pathogenicity traits.

Several microbial plant pathogens colonize xylem vessels including strains of *Clavibacter, Erwinia, Pseudomonas, Ralstonia* and *Xanthomonas* genus and *Xylella fastidiosa* (De La Fuente et al. 2022; Yadeta et J. Thomma 2013). Xylem is composed of pipes transporting water and nutrients from roots to leaves thanks to evapotranspiration process *via* leaf stomata (De La Fuente et al. 2022). Xylem sap is principally composed of water and ions, as well as few amino acids like L-glutamine (Gerlin et al. 2021). Sugars and dioxygen are present in low concentration. Xylem specific pathogens infect and colonize plants by entering *via* wound or natural opening, from leaf or roots. They mainly cause symptoms by obstructing vascular vessels due to cell proliferation, biofilm formation or activation of plant defense (De La Fuente et al. 2022; Yadeta et J. Thomma 2013; Bae et al. 2015). Only a few studies have been conducted to investigate the similarities or differences in the traits that such pathogens utilise to infect the xylem, and the ways in which the constraints within this specific ecological habitat influence the evolution of their infection strategies.

Among xylem colonizing bacterial pathogens, *Ralstonia solanacearum* (*Rs*) is one of the most devastating pathogens (Lowe-Power et al. 2018; Mansfield et al. 2012). *Rs* is a proteobacteria responsible for the bacterial wilt disease affecting more than 250 plant species such as tobacco, potato and tomato (García et al. 2019; Peeters et al. 2013). *Rs* lives in the soil and infects its host plant through its roots. Once inside the roots, *Rs* is transported by the xylem flow up to the leaves. After a few days in, the xylem vessels become blocked, causing the plant to die (Ingel et al. 2022). *Xanthomonas campestris pv. campestris* (*Xcc*) is the causing agent of the black rot disease. This phytopathogen infects especially Brassicaceae vegetables (An et al. 2020; Liu et al. 2022; Vicente et Holub 2013). *Xcc* enters plant through the hydathodes of leaves, which can be recognized by the V-shaped pattern of infection on the leaves (Vicente et Holub 2013). As *Xcc* infects *via* the leaves, it needs to develop strategies to move and colonize against the upward xylem flow. Unlike *Rs, Xcc* takes several weeks to colonize and infect leaves.

*Rs* and *Xcc* deploy a wide arsenal of virulence factors. Some of these infection mechanisms are shared. During the colonization of the xylem, pathogen first needs to adhere to the plant cell *via* pili and adhesins. Both pathogens use the Type II Secretion System (T2SS) to lyse plant cells and obtain nutrients for growth (Alvarez-Martinez et al. 2021; An et Zhang 2024). Another common secretion system used is the Type III Secretion System (T3SS), which secretes effectors (T3Es) that hijack plant immunity (Alvarez-Martinez et al. 2021; An et Zhang 2024). The secretion of exopolysaccharides (EPS) is essential for forming biofilm and increasing the viscosity of the surrounding environment, thereby enabling bacterial cells to remain in the xylem for longer by blocking or limiting sap flow (Bianco et al. 2016; An et Zhang 2024). In *Xcc*, two types of EPS are produced. At low cellular density, activation of *xag* genes leads to the secretion of EPS responsible for cell aggregation. The endomannase enzyme ManA is synthesized and secreted at high cell densities, where it degrades EPS related to *xag* genes leading to the destruction of biofilms. *Xcc* secretes another type of EPS, called xanthan, which is promoted by *gum* genes responsible for an increase in medium viscosity. Finally, swimming and swarming motility are the mechanisms used for moving into and inside the xylem (De La Fuente et al. 2022).

Both *Rs* and *Xcc* rely on complex virulence regulatory networks (VRN) to activate common virulence functions adequately. Indeed, the virulence program is among other mediated by quorum sensing (QS) in both *Xcc* and *Rs* (De La Fuente et al. 2022). QS is a mechanism of cell-to-cell communication that regulates gene expression and cell phenotype in response to external stimuli. The activation of QS signaling pathway is mainly modulated by an increase in cell density, thus augmenting the production of QS signal molecules. Actors of this mechanism are known to regulate the expression of virulence genes and monitor the lifestyle switch in both *Rs* and *Xcc*, despite the fact that they use different signal molecules and regulators : 3-hydroxypalmitate (3-OH PAME) and *phc* in the case of *Rs* (Tarighi et Taheri 2011; Kai 2023; Genin et Denny 2012), and DSF and *rpf* in the case of *Xcc* (He et al. 2007; He et Zhang 2008; Tarighi et Taheri 2011).

The execution of regulatory programs can be difficult to study and resolve, especially in complex environments. Systems biology approaches can be powerful tools for deciphering how complex regulatory programs emerge in response to constraints encountered within the host (Peyraud et al. 2017). We first reconstructed and validated the VRN of *Xcc*. We then used the unique reconstructed mathematical model of the VRN of *Rs* to compare how its regulatory program balances pathogenicity trade-offs with those of *Xcc*, which colonizes the same plant xylem niche. Despite sharing similar pathogenicity traits, we predicted that differences in the virulence regulatory programs would reflect adaptive responses to optimize spread from natural infection niches under the xylem flow constraints.

## Materials and Methods

### Regulatory network reconstruction

The regulatory network of *Xcc* is based on multistate logical modeling. In this model, the state of each entity is computed according to an activation rule that depends on the states of other entities in the network, reflecting the behavior of signaling cascades in biochemical networks. The regulatory network was reconstructed and manually curated based on genome information and integration of literature knowledge from 32 articles, like transcriptomic and phenotypic (**see Supplementary material 4**). The network focuses on molecular and genetic regulations involved in virulence processes: (i) adhesion and aggregation *via* the Type IV pilus and biofilm, (ii) production of viscous exopolysaccharides (EPS), (iii) swimming motility including chemotaxis and flagellum, (iv) plant cell wall (PCW) degradation with the Type II Secretion System (T2SS) and extracellular enzymes biosynthesis and secretion, (v) plant immunity modulation with the Type III Secretion System (T3SS) and T3 effectors. These functions are modulated by environmental stimuli and especially QS. The network is reconstructed based on a methodology developed for *Rs* (Peyraud et al. 2018). Briefly, each interaction is defined by one regulator and one target entity. The type and mechanism of the interaction define the link between the regulator and the target, allowing the rules for target activation to be established. The model is available on an excel and a sbml-qual file (**Supplementary data 1 and 2**). The *Xcc* regulatory network generated in this work was compared to the *Rs* regulatory network model (Peyraud et al. 2018), which is available in **Supplementary data 3**.

### Regulatory steady-state analysis (RSA) simulation

The simulation was done using Regulatory Steady-state Analysis (RSA) (Marmiesse et al. 2015) as described in Peyraud et al 2018 (Peyraud et al. 2018). RSA is a method that allows the first attractor of the system to be found at steady-state by defining the initial state of each entity. Network simulations of the infection mechanism at low and high cellular densities were initiated using data on the composition of tomato xylem sap (Peyraud et al. 2018). For *Xcc* and *Rs*, high cellular density is modeled by setting the concentration of QS inducer, Diffusible Signal Factor (DSF) or 3-OHPAME respectively, above the perception threshold.

### Model performance evaluation

#### Gene expression validation

The performance of *Xcc* regulatory network was evaluated using four transcriptomic data sets (**Supplementary data 4**). Expression level of 113 genes of the network was compared to RNA-seq measurements performed in the following media: NYG (Liu et al. 2013), MOKA (Luneau et al. 2022) and MMX (Liu et al. 2013; Jiang et al. 2018). Gene expression was quantified through reads per kilobase of transcript per million mapped reads (RPKM). RPKM values were converted into binary expression 0/1 by applying a threshold of 75. This threshold was determined based on the RPKM expression value of *hrp* and *xop* genes into the HrpG* condition (Luneau et al. 2022). Constitutive activation of HrpG leads to the activation of the transcription of these genes (Monnens et al. 2024). This allows to determine a threshold that separates genes that are considered expressed from those that are not. The same threshold was applied for the data sets of Liu et al (Liu et al. 2013) and Jiang et al (Jiang et al. 2018).

Accuracy was calculated for each data set:

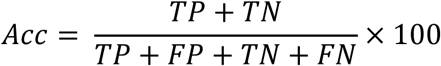

True Positive (TP) corresponds to the number of entities with a simulated and experimentally active expression (1). On the contrary, True Negative is the number of entities that are simulated and experimentally inactive (0). False Positives (FP) and False Negatives (FN) represent discrepancies between simulated and experimental expression levels. FPs occur when the simulation predicts no expression (0), but while expression is observed experimentally (1). FNs occur in the opposite case.

#### Phenotype mutant validation

Phenotypic data from 15 studies (**Supplementary data 5**) collected from different media and growth phase were used to validate the predicted phenotypes of the model. The phenotypes are related to cell adhesion and aggregation (39 conditions), viscous EPS secretion (37 conditions), plant cell wall degradation (7 conditions), and swimming motility (25 conditions) (**Supplementary data 5**). The quantitative experimental results were converted into binary results (activated or not active) and then compared to the binary simulated results. To simulate *in silico* the phenotype of loss-of-function mutant (*i.e.* knock-out gene), the status of the mutated gene was forced to stay null (0) throughout the entire RSA simulation. Phenotype outputs of the knock-out simulation (RSA) were compared to the experimental data.

#### Bacterial spread model

The bacterial spread model presented in this work is based on the regulation program of *Xcc* and *Rs*, as well as the direction of xylem sap flow. Root infection produces forward flow (with bacterial swimming), while leaf infection generates backward flow (against swimming). This model focuses on the expression of swimming phenotype and on production of viscous EPS mediated by quorum sensing. Activation of phenotypic traits was determined based on cell concentration threshold (**Supplementary material** 1). The way in which the xylem exerts its flow affects how cells advance through the stem. The concentration of EPS modifies the viscosity of the xylem sap and then impacts the growth rate of swimming cells (**Supplementary material 2 and 3**). Excess of EPS leads to blockages in the stem, which is modeled by barriers that stop the flow of xylem sap.

The algorithm calculates the cell and EPS concentration based on xylem sap flow and regulation mechanism (**Supplementary figure 1**). Bacterial concentration at each time and distance is calculated as follows:

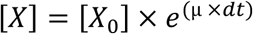

The growth rate (µ) changes depending on the flow and the phenotype of the cell (**Supplementary material 1**). EPS concentration produced by cells was calculated based on the secretion flux of *Xcc* and *Rs*.

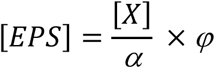

The EPS concentration [𝐸𝑃𝑆] in mmol/L was calculated using the cellular concentration [𝑋] in cell·l^-1^ multiply by α, the number of cells into 1 gDW which is in the order of magnitude of 2.5 x 10^-12^ (Peyraud et al. 2016)) and by φ, the exopolysaccharide secretion flux in mmol/gDW/h.

The impact of EPS concentration on fluid viscosity and ATP cost was estimated through xanthan viscosity (Gao et al. 2023; Mohsin et al. 2021) and for *R.solanacearum* EPS (Drigues et al. 1985) (**Supplementary material 3**).

The impact of motility on bacterial growth rate was assessed through the estimation of ATP cost in mmol/gDW/h (**Supplementary material 2**). This ATP cost is derived from an estimation of flagellar motility in *Escherichia coli* (Schavemaker et Lynch 2022). In addition, Stoke’s law was used to calculate the force required for movement in a fluid (**Supplementary material 2**).

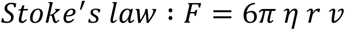

The force 𝐹 (in N) is the product of the viscosity 𝜂 (in Pa.s), the radius of the sphere 𝑟 (in m), and the velocity 𝑣 (in m/s).

Parameters for each of the four models are described in **Supplementary material 1**.

## Results

### Reconstruction of *Xanthomonas campestris* regulatory network

To decipher how the infection program mobilized by *Xanthomonas campestris* within the xylem vessels emerged from the molecular level, we reconstructed the *Xcc* virulence regulatory network. We collected and integrated knowledge about regulation of molecular components from 32 publications (**Supplementary material 4**). This manually curated model contains functional signaling pathways that respond to 43 stimuli such as environmental nutrients, pH, dioxygen, plant stimuli, and QS inducer. These stimuli trigger signaling pathways that regulate the expression of 151 genes encoding virulent factors such as the T3SS and effectors, motility, viscous exopolysaccharides (EPS) production and adhesion (**Figure 1A**). The regulatory network reconstruction resulted in 707 interactions between 478 entities (**Supplementary data 2**). Entities represent genes, RNAs, proteins, but also proteic complexes, metabolites, metabolic reactions, or physical parameters (**Figure 1A**). The QS signaling pathway has been identified as a key hub of the network due to its involvement in 105 interactions that directly regulate five main virulence functions: aggregation and adhesion, viscous EPS production, swimming motility, plant cell wall (PCW) degradation, and modulation of plant immunity. Environmental nutrients, such as amino acids found in rich-media, are tightly connected to motility and modulation of plant immunity (**Figure 1.B**).

**Figure 1.**
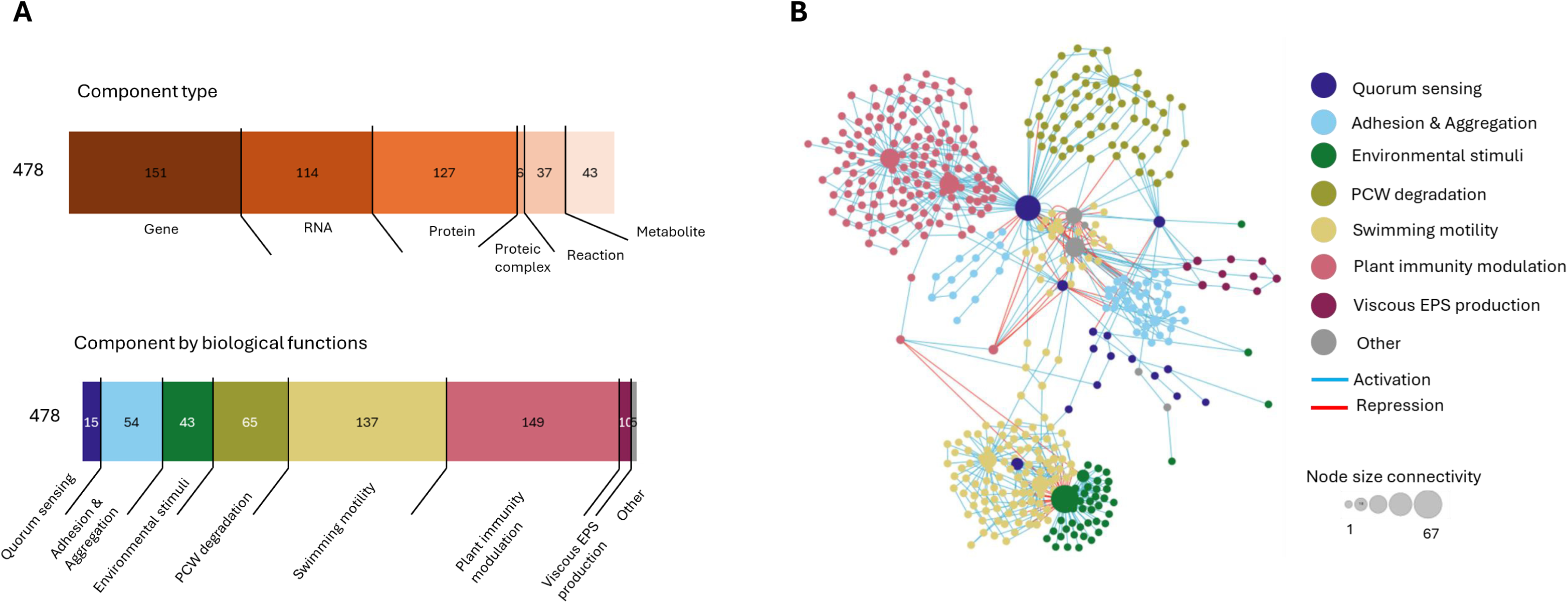
Features of *Xanthomonas campestris pv. campestris* regulatory network model reconstructed. (**A**) Description and metrics of the regulatory network components by molecular type and assignment of their roles in biological functions (**B**) Graphical view of the network components and their interactions colored by their main biological functions.

The reliability of the *Xcc* regulatory network to predict gene expression reprogramming was evaluated by comparing predicted gene expression using Regulatory Steady-State Analysis (RSA) (Marmiesse et al. 2015) and data from four RNA-seq data sets of *Xcc* 8004 (**see Materials and Methods**). The overall accuracy was 71% (**Figure 2A**) with 75% and 76% for NYG and MOKA media (Luneau et al. 2022; Liu et al. 2013) and 70% and 64% for both datasets in MMX medium (Liu et al. 2013; Jiang et al. 2018) (**Supplementary data 4**). More specifically, the network presented a high accuracy to predict expression of genes involved in adhesion and aggregation (99%), EPS production (83%), swimming motility (75%), and modulation of plant immunity (81%). The network performed well in only half of the cases for the expression of genes involved in PCW degradation system, with an average accuracy of 53% (**Figure 2.A**).

**Figure 2.**
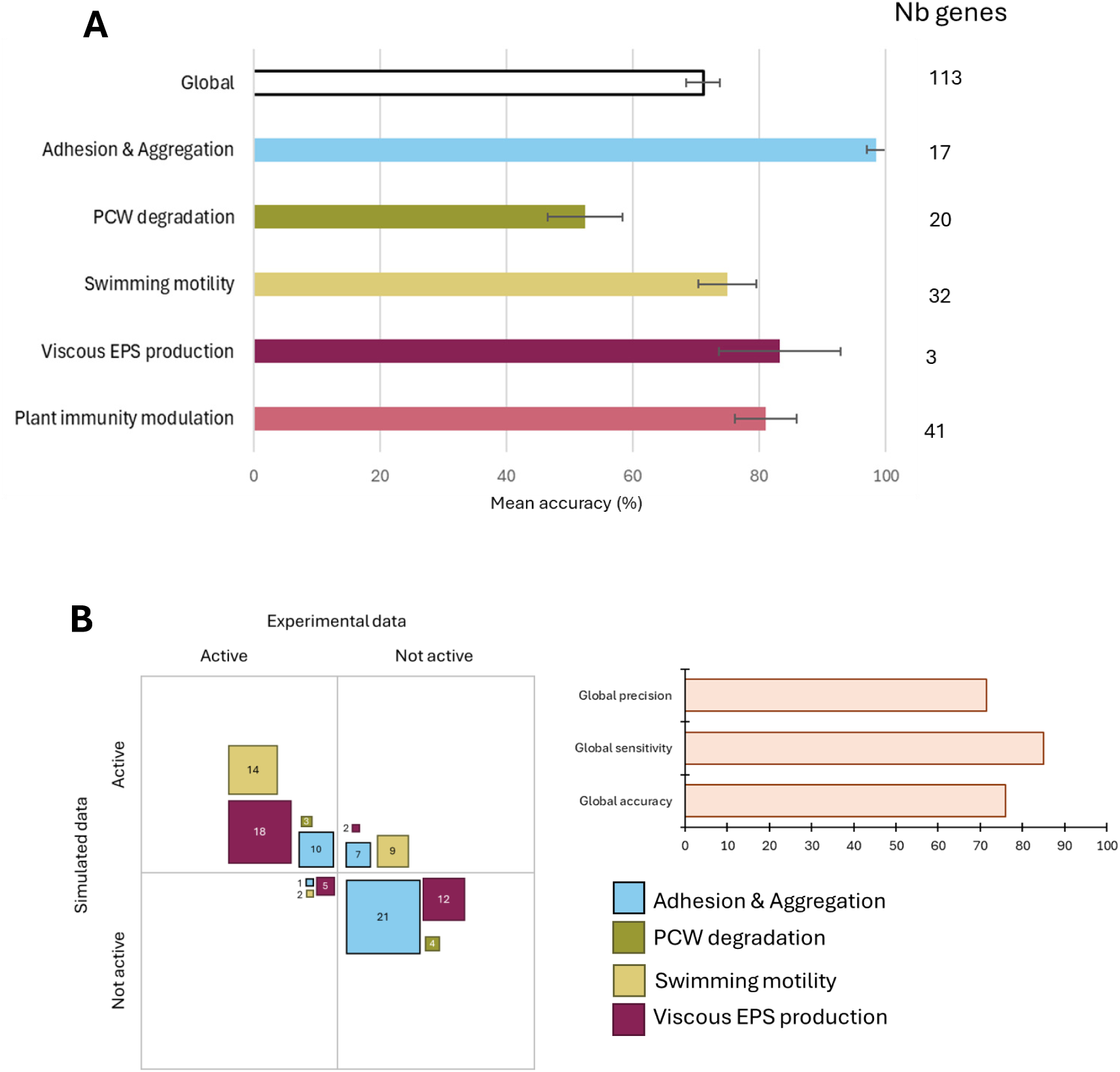
Prediction performance of *Xanthomonas campestris* pv. *campestris* regulatory network model. (**A**) Global and biological functions gene expression accuracy between model simulations and RNA-seq data sets (Luneau et al. 2022; Liu et al. 2013; Jiang et al. 2018) (**Supplementary data 4**) (**B**) Comparison of the model’s prediction of 113 mutant phenotypes and experimental data (**Supplementary data 5**) for four phenotypic traits.

In addition to validating the prediction capacity of gene expression, the activation and repression of phenotypic traits prediction capacity were validated. Simulations with RSA mimicking *in silico* knock-out (KO) were compared with the observed experimental phenotypes of 108 *Xcc* 8004 and XC1 mutants (**Supplementary data 5**). Four global phenotypes were evaluated: aggregation & adhesion, viscous EPS production, swimming motility, and PCW degradation (**Figure 2B**). The model had an accuracy of 76% with 85% precision and 71% sensitivity. Phenotypes of adhesion and aggregation, viscous EPS production, secretion of PCW degrading enzyme were well predicted by the model. In contrast the accuracy for flagellum motility was only at 56% (**Supplementary data 5**).

### *Xanthomonas campestris* pv*. campestris* and *Ralstonia solanacearum* harbor similar virulence related functions coded by unrelated genes

Despite being phylogenetically distant, *Xcc* and *Rs,* both notorious to infect xylem. We therefore investigated whether their virulence regulatory networks had converged to control identical traits using similar genes. Five biological functions were found to be common to both models (**Figure 3**) representing a set of 365 genes, of which only 18% were shared by both pathogens. Structural genes involved in the biogenesis of protein complexes including flagellum, T2SS, T3SS and TIV pilus were overall conserved between *Xcc* and *Rs* with 70% of orthologs genes. In contrast, only three T3Es were orthologs between the two plant pathogens (**Figure 3** **and Supplementary data 6**). In contrast, identical virulence-related functions were performed by totally different sets of genes. As already known, the QS signaling pathway is encoded by two unrelated gene clusters. Moreover secretion of viscous EPS process involves non orthologous genes: *gum* for *Xcc* and *eps* for *Rs*.

**Figure 3.**
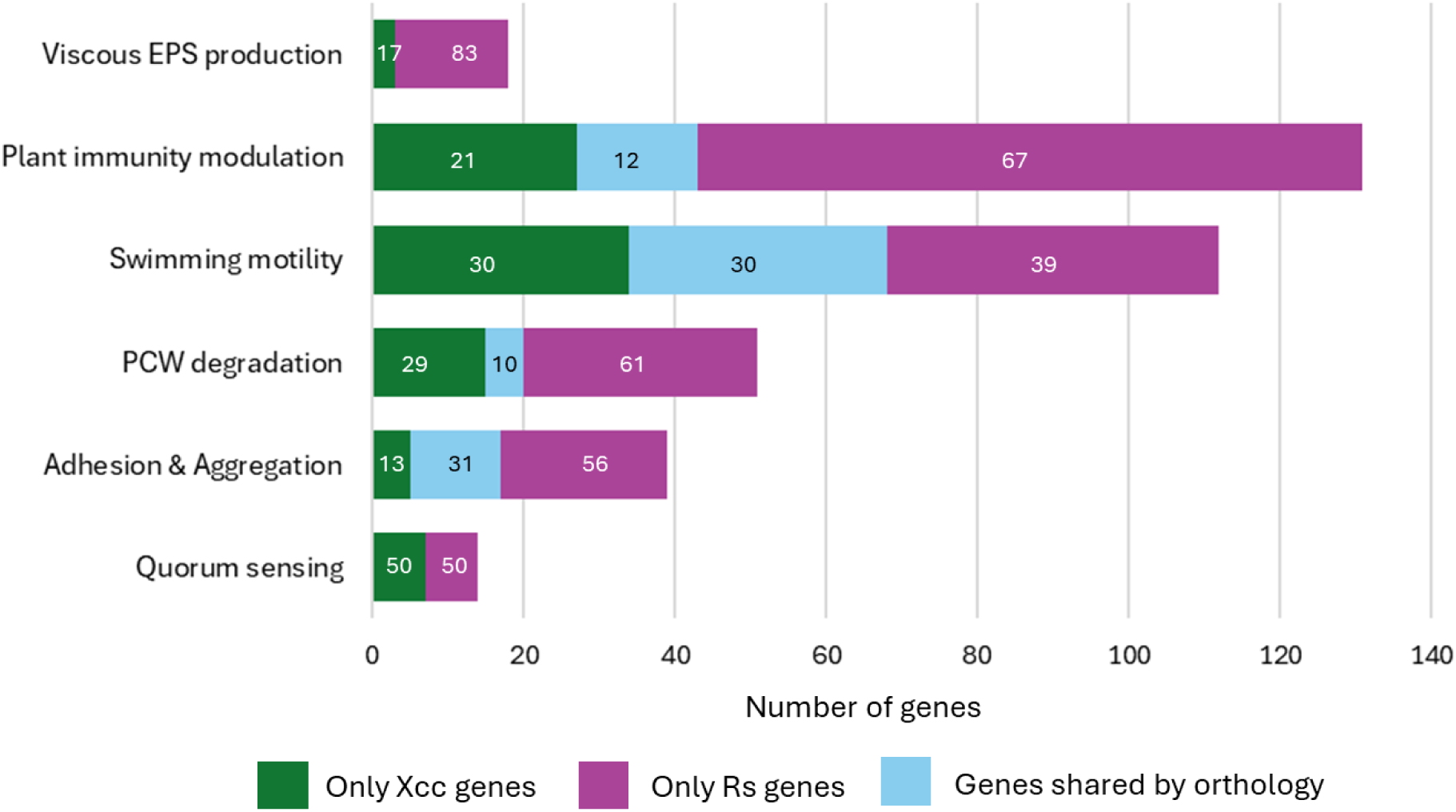
Comparative analysis between *Rs* and *Xcc* for their virulence regulatory network genes, which are assigned to similar biological functions.

Thus, although the virulence functions are identical, some of them are carried out by completely different gene sequences.

### The two xylem infecting pathogens harbor distinct control of viscous EPS and swimming motility

To investigate if the virulence regulatory network of both pathogens converged to a similar regulation of the five common biological functions during xylem colonization, we predicted their regulation at low and high cellular densities (**Figure 4**, see **Supplementary data 7** for model parametrization). *Xcc* and *Rs* execute virulent functions similarly like adhesion and aggregation, in tomato xylem sap. These functions are predicted to be activated at low cell density and not at high cell density in xylem. Furthermore, the modulation of plant immunity, which involvesT3SS and T3Es is predicted to be activated at both low and high population concentrations for *Xcc* and *Rs*. It is interesting to note that three out of the five functions were predicted to be regulated differently including: PCW degradation, swimming motility, and viscous EPS secretion. While *Rs* switches between swimming motility at low density and viscous EPS production at high density, *Xcc* activates EPS secretion independently of the cellular density. *Xcc* swimming motility activation occurs only at high cell density. Concerning PCW degradation, the simulation predicted that *Rs* permanently induces this function, whereas *Xcc* only activates it at high cellular density.

**Figure 4.**
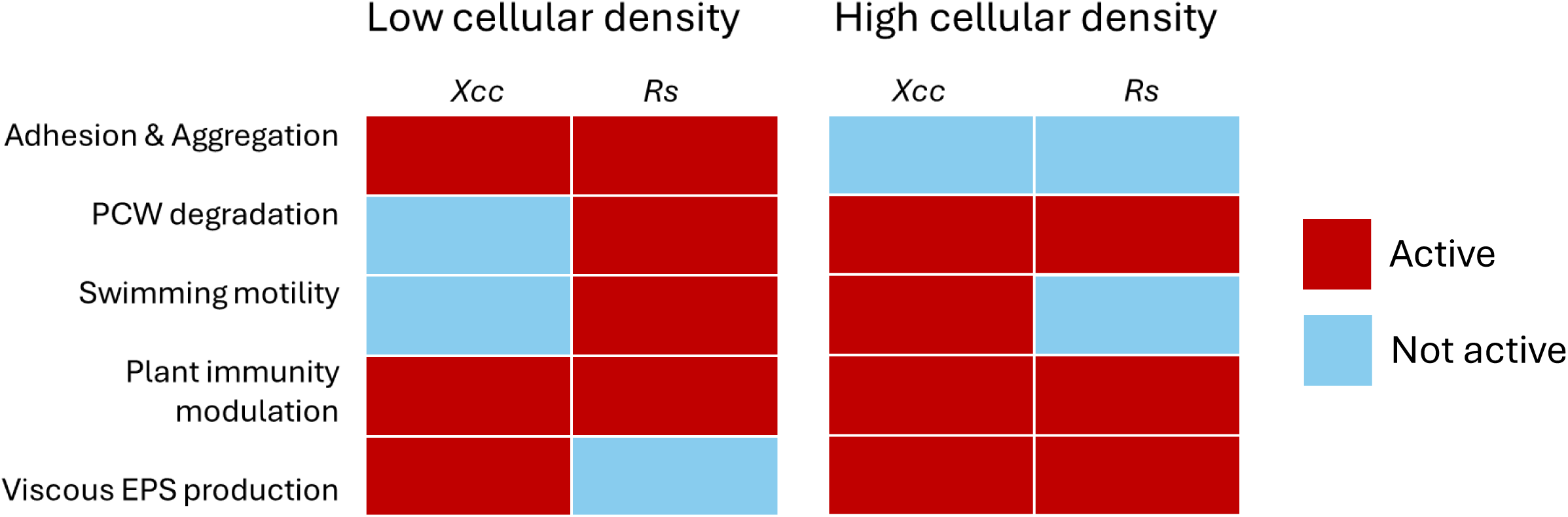
Simulations of five virulence functions status (activated or not activated) of both pathogens which are under the control of the virulence regulatory network. Simulations are performed in xylem sap at low or high cell density, *i.e.* with a lower or higher concentration of QS perception.

Despite having similar virulence functions, the execution of PCW degradation, viscous EPS secretion and swimming motility in the xylem varies between pathogens. Hence, the opposite regulation of EPS secretion and motility observed in *Rs* may be a means to mitigate the trade-off between increasing the viscosity of xylem sap to block the vessel and spread thanks to motility. However, the observation that *Xcc* does not seem to mitigate this trade-off calls into question whether the increase in viscosity caused by viscous EPS may represent a significant cost to the cell (**Figure 5C**).

**Figure 5.**
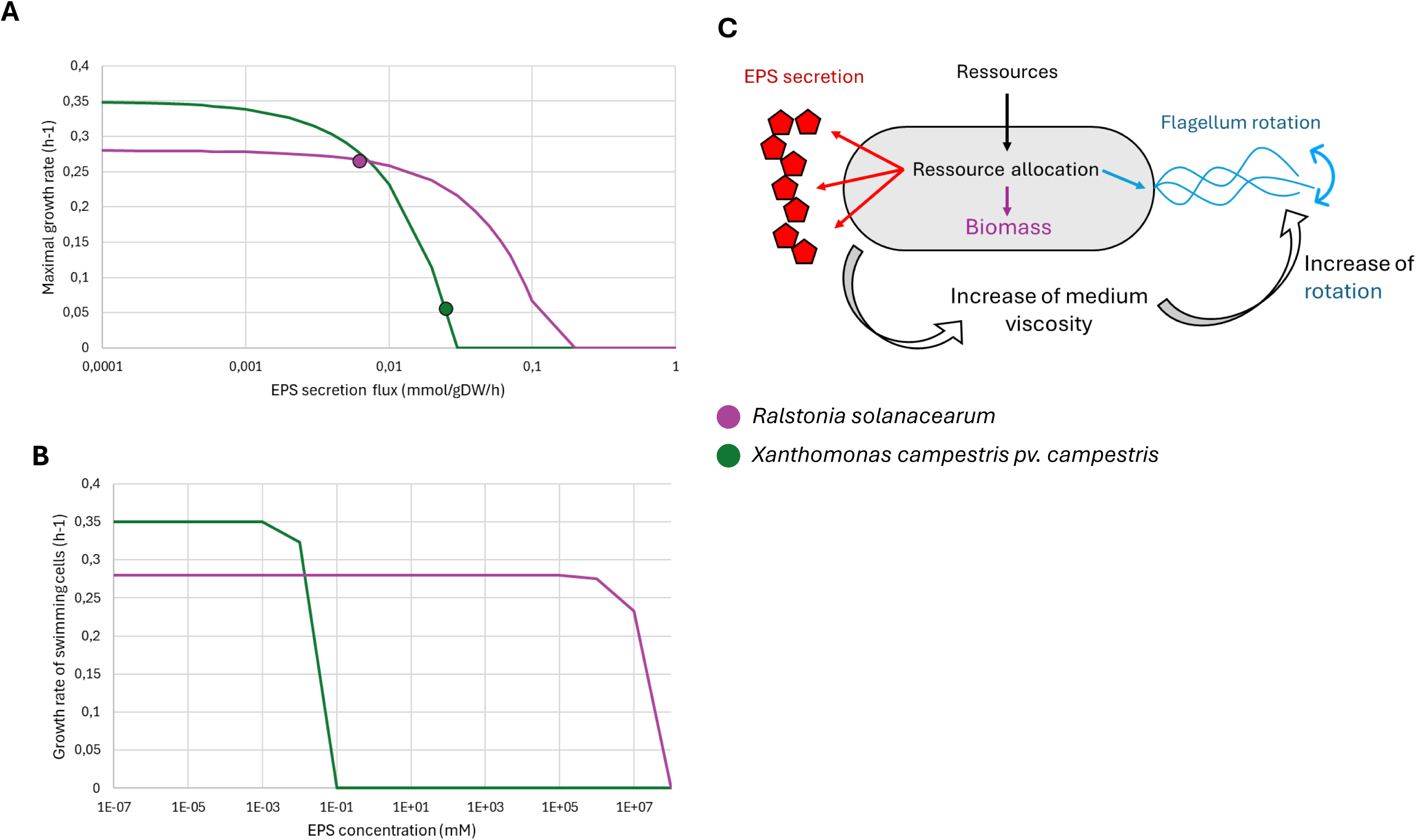
Simulation of the trade-offs between viscous EPS secretion on proliferation and motility for *Rs* and *Xcc.* (**A**) Impact of EPS secretion rate on the maximal growth rate of bacteria due to constraints on resource allocation. (**B**) Impact of EPS concentration in the medium, on the maximal growth rate of swimming cells due to an increase in viscosity, assuming a stable velocity of the cell in the medium. (**C**) Schematic representation of the resource allocation trade-off between viscous EPS secretion, medium viscosity, swimming motility and proliferation. (**A and B**) Line corresponds to simulation results and dot to experimental data.

To investigate the cost of swimming in a viscous medium we built a biophysical model to calculate the cost of a cell moving through a fluid. The model was parametrized using experimental data on the viscosity changes of *Rs* and *Xcc,* as well as their respective EPS and motion speed measurements (**see Supplementary material 1**). EPS production impacts differently *Xcc* and *Rs* (**Figure 5A**). Indeed, the biosynthesis and secretion cost of xanthan much more severely penalizes the proliferation rate of *Xcc* compared to *Rs*. The model predicted that motility in non-viscous fluid does not significantly impact the growth rate of either *Rs* and *Xcc* (**Supplementary material 1**) due to negligible friction force in the case of weak xylem sap flow and cell velocity. Increase in the viscosity leads to more energy deployed to swim inside xylem sap and thus decreases the growth rate (**Figure 5.B**). At the same EPS concentration,

xanthan results in a much more viscous medium (**Supplementary data 9**). This results in a very rapid decrease in the growth rate of swimming cells at 10^-3^ mM EPS. For *Rs,* a higher EPS concentration is required to impact the growth rate: 10^6^ mM.

### The regulatory program of the pathogens is adapted to the constraints generated by their infection site and xylem flow directionality

Swimming motility and secretion of viscous EPS may impact dispersal within the xylem and proliferation capacity of the pathogens (Peyraud et al. 2016). Hence, contrasted strategies employed by each pathogen to mobilize these functions may respond to specific constrains during xylem colonization. To investigated how the regulatory program impacts xylem colonization, we developed a bacterial spread mathematical model within this habitat.

The mathematical model mimics the colonization and spread of the bacterial populations within the xylem vessel. It simulates the following : (i) the speed of the bacterial cells within the xylem vessel, depending on their intrinsic velocity and constraints when swimming forwards or backwards in xylem sap composed of flow and viscosity; (ii) the proliferation rate of the cells depending on the cost of production of their virulence functions; (iii) manipulation of the xylem flow rate and viscosity through the secretion of viscous EPS and aggregation (**see Material and Methods and Supplementary material 1** for model details and parametrization).

Colonization efficiency of xylem vessels by both pathogens was first compared from root infection, *i.e* the pathogens spread within the xylem in forwards direction of the flow (**Figure 6.A**). Under these conditions *Rs* is predicted to reach higher cell density in few days, and also spread up to 100 mm in 6 days. In contrast *Xcc* does not spread due to the prediction of early establishment of a biofilm due to activation of aggregation and viscous EPS secretion, which ultimately stops xylem flow. In the case of xylem colonization *via* the leaves, *i.e.* the pathogens spread backwards (**Figure 6.B**), *Rs* does not spread since the velocity of the flow used exceeds the motility velocity of the cells (**Supplementary material 1**). *Xcc* first forms barriers through aggregation and production of viscous EPS, which blocks the xylem flow. This allows the bacterial populations to spread backwards up to 25 mm within 20 days.

**Figure 6.**
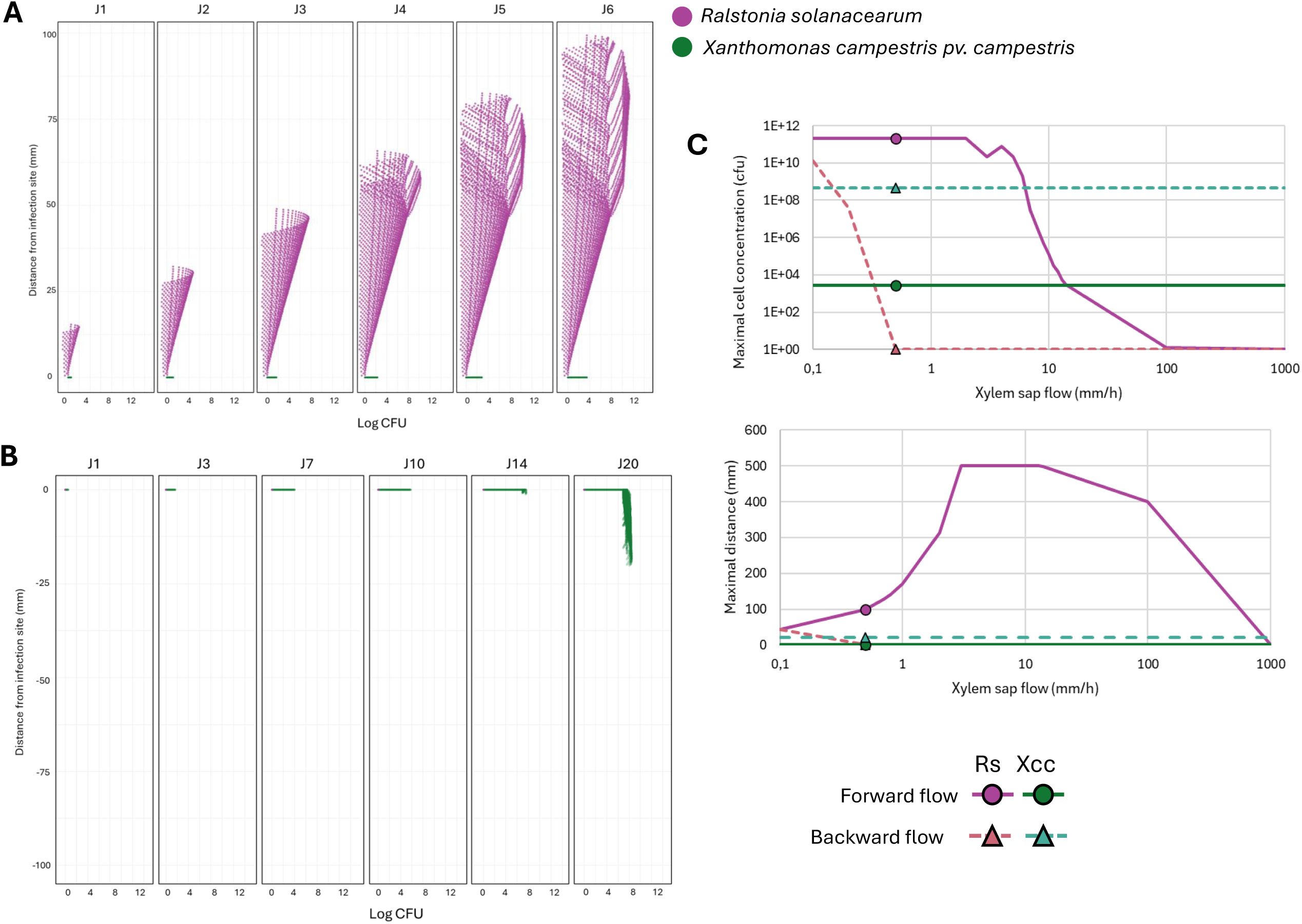
Bacterial dynamic spread simulation of xylem infection of *Rs* (purple) and *Xcc* (green). (**A**) Dispersion model simulation from root, *i.e.* in the direction of the xylem flow (**B**) Dispersion model simulation from leaf, in the opposite direction to the flow of xylem sap (**C**) Sensitivity analysis of bacterial colonization performance, bacterial population (upper panel) and distance spread (lower panel), under the constraints of flow of xylem sap. Line corresponds to simulation results and dot to experimental data.

We conducted a sensitivity analysis to identify the parameters shaping the performance of both pathogens in infecting the xylem (**Supplementary data 8**). Among the parameters used, the xylem sap flow parameter was found to be the main environmental constraint. Varying the xylem sap velocity from 0.1 to 1000 mm/h showed that only *Rs* infection is significantly impacted (**Figure 6.C and Supplementary data 8**). Regardless of the strength of xylem sap flow, neither the maximal cell concentration nor the maximal distance of *Xcc* are affected by this parameter. This demonstrates that the *Xcc* regulatory program enables robust infection of plant xylem, even under high sap flow pressure. However, this robust infection is at the expense of a slow speed of infection, around 20 days. Conversely, *Rs* spread is highly sensitive to the xylem sap velocity. Indeed, in forward flow, an increase in sap speed of up to 2 mm/h leads to an increase in the maximum colonization distance but does not affect the maximum cell concentration. The analysis shows a stagnation from 3 mm/h to 13 mm/h due to the maximum distance calculated by the model at 500 mm. Under backward flow, only a sap velocity lower than 0.2 mm/h allows *Rs* to spread. This speed is an order of magnitude lower than the measured bacterial speed (**Supplementary material 1**), and Rs motility could be effective for sap flow at this level.

Hence, modeling of *Rs* and *Xcc* infection strategies based on swimming motility, viscous EPS secretion, and aggregation - all three of which are controlled by QS - demonstrated that the regulation of these functions is adapted to their natural infection site, with contrasted robustness towards the strength of xylem sap flow.

## Discussion

In this study we used systems biology approaches to decipher the causal mechanisms underlying the trade-offs between pathogenicity traits. We modelled the physical constraints, such as the impact of viscosity on cellular motion in a fluid, as well as thermodynamic constraints on cell proliferation, such as limited energy and matter allocation to pay metabolic costs. The different proliferation and invasion velocities observed between *Rs* and *Xcc* (**Figure 6**) could be partially explained by the high cost of viscous EPS secretion in *Xcc*, which reflects a trade-off between EPS production and proliferation. In addition, the regulatory program of *Rs* mitigates the trade-off between swimming motility and the increased viscosity of xylem sap caused by the secretion of viscous EPS.

For *R*s it has been already demonstrated *in silico* and experimentally that activations of EPS secretion induces a significant decrease in the growth rate of cells, and this trade-off arises from constraints in the allocation of resources (Peyraud et al. 2016). Calculation of the impact of EPS secretion on growth in this work show a 7% decrease in the growth rate for *Rs,* which is similar to the results obtained by Peyraud et al. (Peyraud et al. 2016) and a 84% decrease for *Xcc* (**Figure 5A**). The cost of xanthan production on *Xcc* growth appears to be a major burden, strongly reducing the colonization speed of the pathogen. Xanthomonads are notorious for their slow growth in culture (Ruissen et al. 1993), which is due to massive production of xanthan gum notably used in industry (Oliveira et al. 2025). The trade-off between xanthan production and proliferation has already been demonstrated *in silico* and experimentally in bioreactor (Schatschneider et al. 2013). The differences between *Rs* and *Xcc* in managing this trade-off are significant: QS activates viscous EPS production in *Rs* at high cell density, leading to rapid proliferation at low cell density. In contrast, *Xcc* produces viscous EPS also at low cell density, which slows down the speed of colonization within the xylem.

The second trade-off modeled in the current work is related to the secretion of viscous EPS and cell motility. The production and biosynthesis of viscous EPS already costly for the organism, leads to an increase of medium viscosity (Cope-Arguello et al. 2026). At the same cell velocity, the cost of motility increases due to greater friction forces in a viscous medium, therefore decreasing the available energy for cell proliferation (**Figure 5B**). Reduction of cell velocity by increasing medium viscosity has been reported for several bacteria, including *Bacillus megaterium*, *Escherichia coli* and *Pseudomonas aeruginosa* (Schneider et Doetsch 1974; Magariyama et Kudo 2002), which display a similar trend to that predicted in this study. Hence, the simultaneous activation of viscous EPS secretion and motility appears to be a sub-optimal regulatory strategy. The regulatory program of *Rs* mitigates well this trade-off and thus enables very fast spread within the xylem in few days (**Figure 6A**) (Xu et al. 2022). Similarly, the opportunistic human pathogen *Pseudomonas aeruginosa* (*Pa*), regulates biofilm production and motility in the same way as *Rs*, planktonic behavior at low cell density, followed by biofilm formation through EPS secretion at high cell density (Carey et al. 2024; Kennelly et al. 2024; Liao et al. 2022). QS appears to be a common signal used to control the trade-off between motility and EPS production to increase medium viscosity or cell attachment (Deep et al. 2011). Interestingly, another xylem colonizing pathogen that is closely related to *Xcc*, *Xylella fastidiosia*, exhibits a similar trade-off between motility and biofilm formation under the control of the same DSF/*rpf* regulatory system as *Xcc* (Portaccio et al. 2025). However, the DSF*/rpf* system in *X. fastidiosa* works differently than *Xcc* due to the additional trade-off constraints on the dispersal capacity of the pathogen within the xylem and its transmission to the insect vector, which requires a high adhesion capacity (Chatterjee et al. 2008).

The results highlight the impact of the bacterial infection program on manipulating xylem flow, which involves increasing in viscosity in addition to forming biofilm. This blocks the xylem flow and then triggers the plant symptoms. Considering only the trade-off between EPS secretion and motility, the strategy deployed by *Rs* appears more optimal, as evidenced by its faster observed colonization and spread within the xylem (**Figure 6**). However, modeling the physical constraints of the xylem flow strength and directionality related to the infection site, whether in the leaves or roots, revealed that *Xcc* regulatory program is well adapted to migration backwards against the xylem flow. Dispersal distance and population dynamics are coherent with the order of magnitude of experimental data for both *Rs* (Baroukh et al. 2025) and *Xcc*. Previous works demonstrate high and fast spread dynamic of Rs during colonization of young tomato plant xylem (Jiang et al. 2016; Xu et al. 2022). The dispersal dynamic of *Xcc* is in line with experimental observation of colonization of Brassicaceae plants (Taks et al. 2025). Therefore, our study suggests that xylem flow is a key parameter in understanding the regulatory program deployed by xylem infecting pathogens.

It has been demonstrated that the fluid flow in vessels may also impact QS signaling by washing out autoinducer molecules (Kim et al. 2016). Interestingly, a study on *P. aeruginosa* discovered the presence of a fluid response sensor (*froR*) that impacts pilus expression and is required for spread in a fluid environment (Sanfilippo et al. 2019). In addition to the flow, viscosity could also impact regulation and gene expression. In *Vibrio cholerae*, an increase in media viscosity induces higher virulence expression (Häse et Mekalanos 1999). For now, no such fluid strength or viscosity sensor has been discovered for *Xcc* and *Rs.* One major driver of variation in xylem sap strength is the day/night cycle (Windt et al. 2006). Regarding the importance of our simulations for the backward and forward migration of the pathogens, such drastic variation between day and night, when evapotranspiration pulls a strong and weak demand for water, should strongly impact the invasion of pathogens. Comparisons between simulation and experiment were performed to study the diurnal variations in xylem sap flow on *X. fastidiosa* biofilm development and spread (Walker et al. 2024). The bacterial spread model developed in this study could provide additional insights by varying the sap flow over time to simulate the diurnal differences in bacterial invasion.

In this study we predicted the regulatory programs deployed by pathogens based on the interaction between molecular components that control genes involved in pathogenic traits. The networks investigated for the two pathogens, as well as those of similar xylem colonizing agents such as *X. fastidiosa*, demonstrate a high degree of modularity in the genes and their biological functions. Indeed, many pathogenic traits can be coded by well conserved genes, such as the T3SS, T4 pilus and flagellum gene clusters (McCann et Guttman 2008; Stroml et Lory, s. d.; Ligthart et al. 2020), or can be performed using totally unrelated genes that perform an identical function, such as the cluster responsible for viscous EPS secretion. Also, the QS function is coded by two different systems in *Rs* and *Xcc*, whereas the components closely related to each other in *Xcc* and *X. fastidiosa* exhibit slight variations that can lead to opposite behavior (Chatterjee et al. 2008). The study supports the idea that genes recruited for pathogenicity are highly modular, and that the regulatory network is highly plastic, allowing to fine-tune the activation of the appropriate pathogenicity determinants. This highlights the importance of considering a system-level perspective when trying to understand and predict the emergence of pathogenicity.

## Conclusion

The *Xcc* regulatory network reconstructed in this work gathered regulatory mechanisms of various pathogenicity traits that are involved in virulence and plant infection. These functions include aggregation and adhesion, viscous EPS production, swimming motility, degradation of the plant cell wall (PCW) and modulation of plant immunity. They are mediated by environmental stimuli and are notably orchestrated by QS signaling pathways. Comparison of *Xcc* VRN with the one from *Rs* revealed strong similarities in the main virulence functions, despite their distant phylogeny. VRN simulations showed differences in the execution of swimming motility, adhesion and aggregation and viscous EPS secretion traits, which are regulated by QS. Construction and simulation of a mathematical model revealed a trade-off between (i) EPS production and cell growth rate and (ii) EPS medium concentration, *i.e.* viscosity, and motility. The differences in the execution of viscous EPS secretion and swimming motility conferred by the regulatory program provide advantages in terms of dispersal and local infection niche. Indeed, simulations of the bacterial spread model show that both *Xcc* and *Rs* colonize their natural infection sites (leaf and root, respectively) more efficiently. While the results presented here are consistent with previous studies in terms of order of magnitude, further validation of the model predictions could be achieved through experimental methods such as using an *in vitro* microfluidic setup to mimic xylem sap flow or *in planta* observation.

## Acknowledgments

No conflict of interest is reported by the authors. We would like to thank J. Luneau for his careful reading of the manuscript and helpful advices. This PhD was funded by ANRT (CIFRE N° 2022/0694). This work was also supported by the French National Research Agency (ANR-20-PCPA-0009).

---

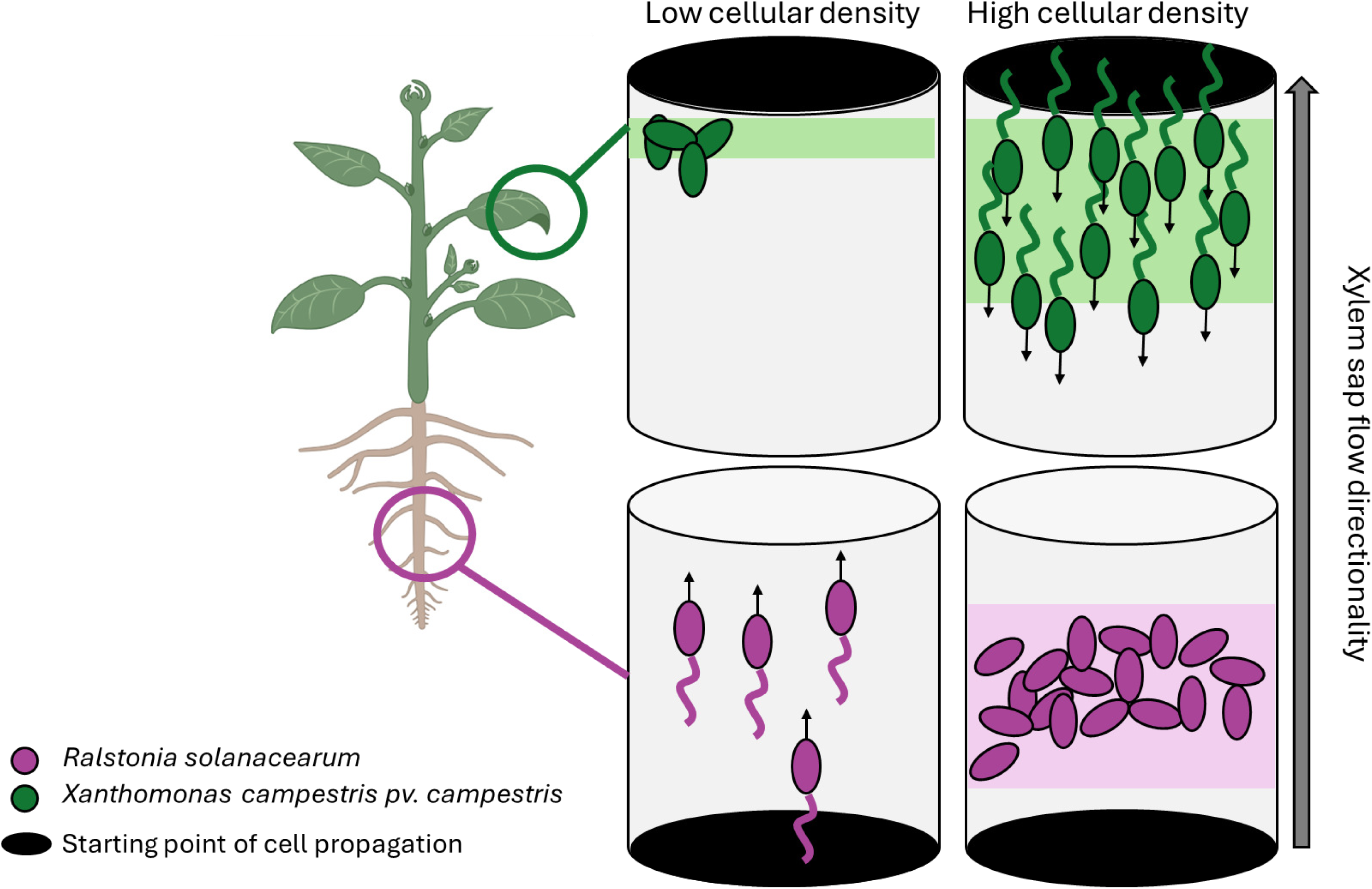

